# Variation in morphology among populations of threespine stickleback (*Gasterosteus aculeatus*) from western Newfoundland, Canada

**DOI:** 10.1101/2022.09.08.507156

**Authors:** RJ Scott, GE Haines, NR Biedak, JA Baker

**Affiliations:** School of Science and the Environment, Grenfell Campus, Memorial University of Newfoundland and Labrador, Corner Brook NL, Canada, A2H 6K8

**Keywords:** anti-predator armor, body shape, geometric morphometrics, geographic variation, ecological speciation

## Abstract

Freshwater populations of threespine stickleback (*Gasterosteus aculeatus*) have diverged from their marine ancestor and show extensive variation among populations throughout most of their range. However, phenotypes of freshwater populations from the east coast of North America does not appear to demonstrate extensive variability observed throughout the rest of the species’ range. The relative young age of east coast North American populations may explain the apparent lack of freshwater variability. On the other hand, populations in this part of the species’ range have not received the level of attention as have populations throughout the rest of its range and the low level of variability observed may simply reflect lack of data for this region. We examined morphological traits (including linear measurements of body armor and body shape) of stickleback from 52 locations representing marine and freshwater populations along the west coast of Newfoundland, Canada. We found that variability in morphological traits among freshwater populations was much more extensive than previously assumed and similar to patterns observed throughout the rest of the species’ range.

## Introduction

Adaptive variation among populations of a species (geographic variation) has been recognized for centuries. Mayr and others envisioned variation of this nature as the starting point for speciation (Liou & Price 1994, Ritchie 2000, Schluter 2000a). Fishes occupying freshwater systems in north temperate regions demonstrate adaptive variability among populations in a number of traits across several families, including salmon, trout, charr, whitefish, sunfish and stickleback, and have contributed to our understanding of the role of microevolutionary processes in producing biodiversity (Robinson and Wilson 1993, Foster et al. 1998, Ehlinger 1999; Losos 2010; Foster 2013; Foster et al. 2015, 2019).

Threespine stickleback (*Gasterosteus aculeatus*, henceforth stickleback) provide a remarkable example of diversification among freshwater populations in several traits, including feeding and anti-predator morphology (e.g. McPhail 1984, Bell et al. 2004; Spoljaric and Reimchen 2007; Barrett 2010; Reimchen et al. 2013), courtship behavior and mate choice (e.g. Foster et al. 2019; Scott 2004), nuptial signals (e.g. Scott 2001, 2011, 2022), life history (e.g. Baker et al. 1998; 2005; Kurz et al. 2016) and physiology (e.g. Guderly et al. 1994; Black et al. 2014; Graham et al. 2018; Scott and Black 2018). Marine stickleback are monomorphic in these traits throughout their range, and represent the ancestral state for all freshwater populations; freshwater populations were established through colonization by marine stickleback following recession of the Wisconsin glaciers roughly 12-15000 y.b.p. (Bell 1987; Bell and Foster 1993; Foster et al. 1998, 2015; Walker and Bell 2000; Hunt et al. 2008; Reid et al. 2021, Fang et al. 2018).

Variability in antipredator armor and body shape among freshwater stickleback populations, and relative to the ancestral marine form, has been studied extensively across much of the species range (e.g. Giles 1983; Campbell 1985; Bell and Orti 1994; Walker 1997; Klepaker and Østbye 2008; Spoljaric and Reimchen 2007). Marine threespine stickleback possess a set of bilateral armor plates (32-34), a robust pelvic girdle (fused ventral plates and bilateral ascending processes), and prominent dorsal and pelvic spines (see Fig. 1 of Bell et al. 2004; Fig. 1 of Klepaker and Østbye 2008; Fig. 2 of Klepaker et al. 2012). Antipredator armor among freshwater populations ranges from individuals possessing the complete sets of lateral armor plates, similar to the marine forms, to reduced plate number (Bell et al. 1993), as well as reduced dorsal and pelvic spine length, and reduced pelvic girdle size. Repeated reductions of armor occurs by different processes, with reduction in plate number tending to occur by transportation of low-plated alleles between freshwater populations via migration through marine environments (Schluter and Conte 2009; Kingman et al. 2021), while pelvic reduction often occurs as a result of *de novo* mutations in the fragile genomic region associated with the *Pitx1* gene (Xie et al. 2019). Some freshwater populations have lost some or all components of their antipredator armor (Reimchen 1980; Edge and Coad 1983; Giles 1983; Campbell 1985; Reimchen et al. 2013).

**Figure 1.**
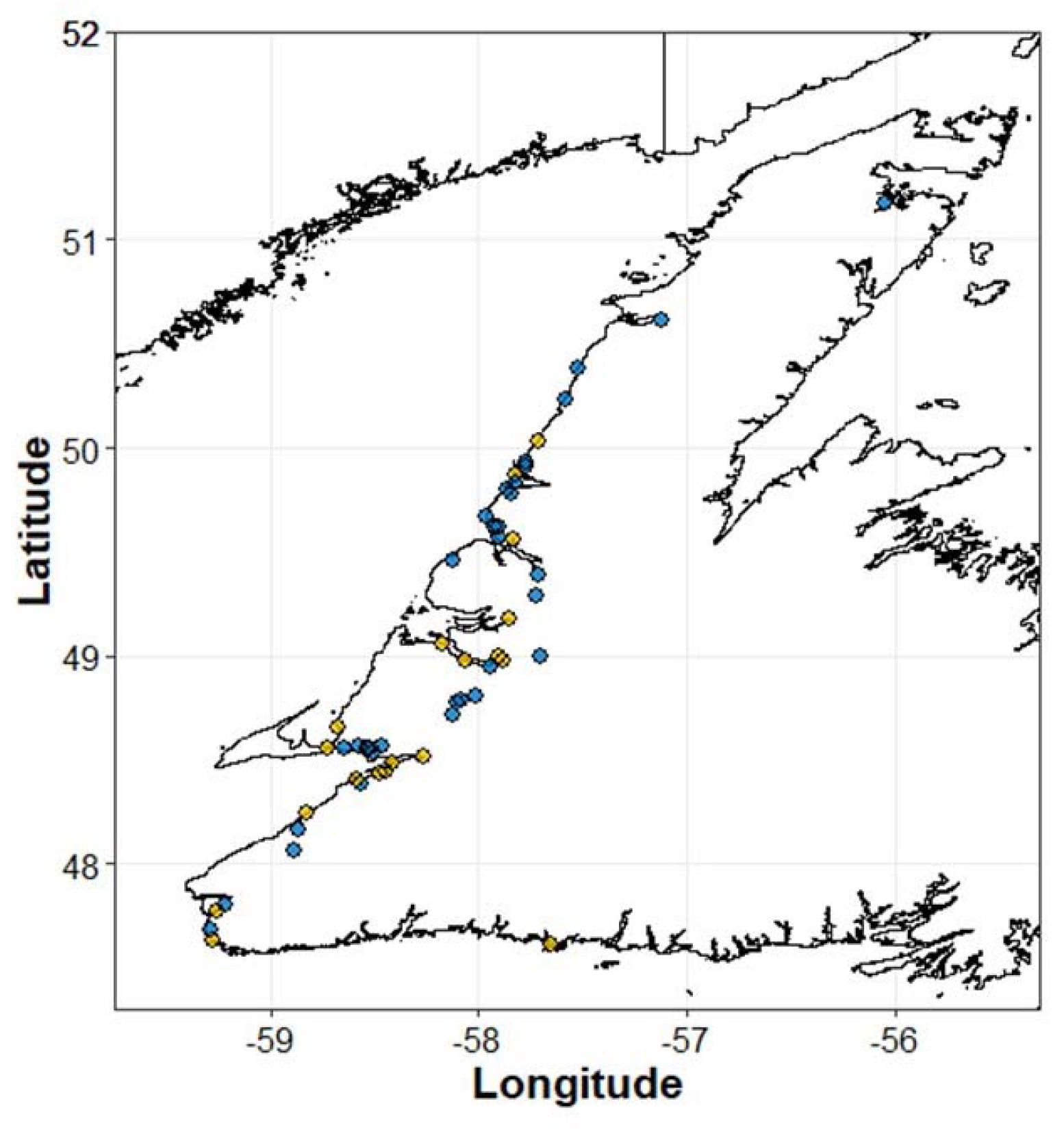
Location of sampling sites from western Newfoundland. Blue (dark) dots indicate freshwater sites and yellow (light) dots indicate marine/estuary sites.

**Figure 2.**
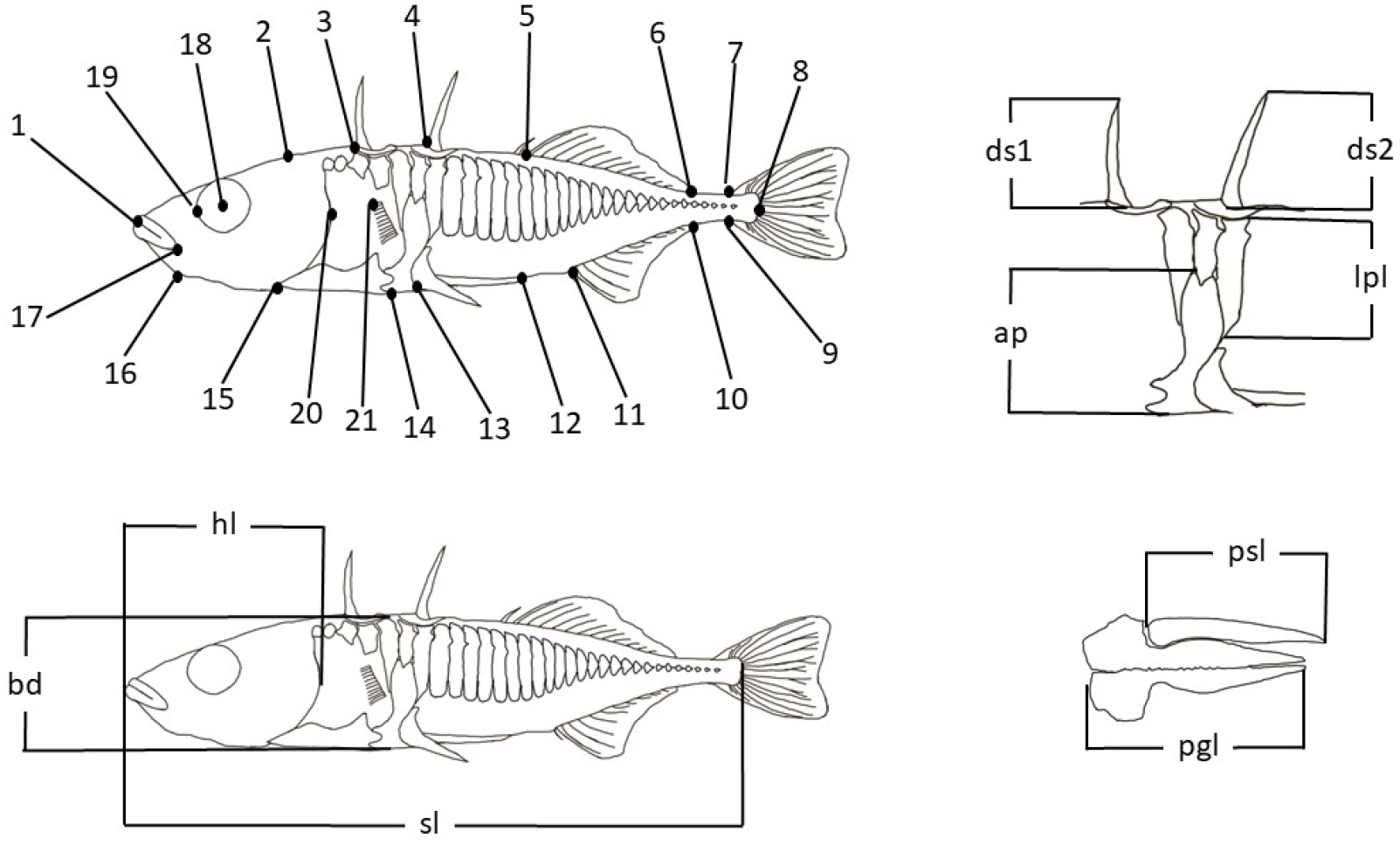
Location of landmarks 1-21 for geometric morphometrics (top left panel) and linear measurements (bottom left – lateral view and top right – lateral view of dorsal spines, ascending process and 3 fused lateral plates magnified) and ventral (bottom right - pelvic plate and spine magnified; sl – standard length; hl – head length; bd – body depth; ds1 – 1^st^ dorsal spine length; ds2 = 2^nd^ dorsal spine length; ap – ascending process length; lpl – lateral plate length; pgl – pelvic girdle length; psl – pelvic spine length).

Body shape varies among lake populations depending on predator presence/absence and specialization on benthic or limnetic food resources (McPhail et al. 1984; Lavin and McPhail 1985; Schluter 2000b; Roy et al. 2010; Willacker et al. 2010; Østbye et al. 2016; Pistore et al.2016). For example, stickleback specializing on planktonic prey tend to have shallower bodies, longer snouts and larger eyes and stickleback specializing on benthic prey tend to have deeper bodies, and smaller eyes both relative to the ancestral marine form (Walker 1997; Walker and Bell 2000). Freshwater populations of stickleback have diverged rapidly and extensively from the marine ancestor in several well-studied regions, including Alaska (Walker 1997; Walker and Bell 2000; Arif et al. 2009; Aguirre and Bell 2012), British Columbia (Bentzen et al. 1984; Lavin and McPhail 1986; Schluter 2000b; Schluter and McPhail 1992; Foster et al. 1998; Vamosi and Schluter 2004), Scandinavia (Klepaker 1996; Klepaker et al. 2012; Klepaker and Østbye 2008; Myhre and Klepaker 2009), the United Kingdom (Coyle et al. 2007; Dingemanse et al. 2007; Ravinet et al. 2013; Klepaker et al 2016; Magalhaes et al. 2016), and Japan (Kakioka et al. 2020; Yamasaki et al. 2019; Ravinet et al. 2021). In each of these regions, the evidence indicates broad-scale phenotypic diversification resulting from natural selection driven by the same environmental factors, particularly features of the habitat and the suite of predators. In contrast, the relatively limited studies on stickleback populations in northeastern North America, especially in Newfoundland and Labrador, suggest limited variation in morphology with relatively minor changes relative to the ancestral marine form (Garside and Hamor 1973; Hagen and Moodie 1982; Coad 1982; Haines 2022). Stickleback populations in eastern North America are the youngest set of stickleback populations spanning such a large portion of the species’s global range (Fang et al. 2018). The low amounts of variability among eastern North American stickleback populations could be due to lack of genetic diversity following at least two historical range expansion events (Fang et al. 2018; Fang et al. 2020). On the other hand, the relative lack of stickleback research on the Atlantic coast of North America may not have captured the level of variability present in this region.

We collected stickleback from freshwater and marine sites along the west coast of insular Newfoundland to examine variability in antipredator morphology and body shape among freshwater populations and between freshwater and marine populations. This study represents the most comprehensive single study of threespine stickleback variability for this number of traits from eastern North America.

## Methods

We collected stickleback from 19 marine/estuarine and 33 freshwater sites (Fig. 1, Table 1). Freshwater sites ranged from small ponds to larger lakes. Physical attributes (lake area, elevation, straight line distance from the ocean, run of river distance from the ocean and average gradient) were estimated using a digital elevation model of the west coast of Newfoundland and the National Hydrology Network in ArcGIS. All sites were under the Laurentide ice sheet during the Wisconsin glaciation. Sites in the southern sample location have been ice-free for approximately 15000 years and those in the north have been ice-free for between 8500 to 9000 years (Liverman 1994), a time interval that corresponds to the spread of threespine stickleback to eastern North America (Fang et al. 2018). We used the information in Liverman (1994) and Daly et al. (2007) to estimate the age (years since glacier free) for each location sampled in this study (Table 1).

**Table 1.** Characteristics of sites sampled for threespine stickleback in western Newfoundland. The three-letter location code (in parentheses for each location name) correspond to the labels in figures throughout the paper. Average gradient was calculated by dividing lake elevation by straight-line distance to the ocean.

| Location | Lat (°N) | Lon (°W) | Lake Area<br>(km<sup>2</sup>) | Elevation<br>(m a.s.l.) | Estimated<br>age (years) | Distance from the<br>ocean |  | Average<br>Gradient |
| --- | --- | --- | --- | --- | --- | --- | --- | --- |
|  |  |  |  |  |  | Straight<br>line (m) | Run of river<br>(m) |  |
| Aband. Limestone Quarry (alq) | 48.380139 | 58.561154 | 0.02 | 56 | 13800 | 3662 | 6527 | 0.250 |
| Estuary at Barachois Brook (bbe) | 48.448011 | 58.441455 | NA | 0 | 13600 | 0 | 0 | 0.000 |
| Bonne Bay Little Pond (bbl) | 49.3899 | 57.7081 | 2.82 | 62 | 10200 | 30061 | 36943 | 0.002 |
| Bennett Lodge Pond (bep) | 50.2269 | 57.5862 | 0.06 | 5 | 8700 | 681 | 996 | 0.007 |
| Berry Hill Pond (bhp) | 49.62339 | 57.9212 | 0.12 | 32 | 9700 | 3023 | 3383 | 0.003 |
| Big Pond (bip) | 47.679079 | 59.292122 | 1.02 | 2 | 14500 | 2714 | 3069 | 0.002 |
| Blue Pond (blp) | 48.7825 | 58.082778 | 0.21 | 187 | 12400 | 41692 | - | 0.005 |
| Bog near Ned's Pond (bog) | 48.557551 | 58.522145 | 0.05 | 69 | 10100 | 4097 | 7040 | 0.017 |
| Barachois River Estuary (bre) | 48.241555 | 58.828369 | NA | 0 | 14000 | 0 | 0 | 0.000 |
| Cook's Brook Estuary (cbe) | 48.974311 | 58.06527 | NA | 0 | 11800 | 0 | 0 | 0.000 |
| Cow Head Pond (chp) | 49.9066 | 57.7757 | 0.27 | 30 | 9100 | 2707 | 3538 | 0.011 |
| Codroy Pond (cop) | 48.061982 | 58.881807 | 1.26 | 140 | 14300 | 42633 | 57869 | 0.003 |
| Cape Ray Estuary (cre) | 47.627218 | 59.281081 | NA | 0 | 14500 | 0 | 0 | 0.000 |
| Dear Arm Estuary (dae) | 49.556972 | 57.832509 | NA | 0 | 9700 | 0 | 0 | 0.000 |
| Downes Pond (dop) | 49.9274 | 57.7712 | 0.07 | 4 | 9100 | 2597 | 4086 | 0.002 |
| Frenchman's Cove (fce) | 49.055947 | 58.175643 | NA | 0 | 11800 | 0 | 0 | 0.000 |
| First Pond (fip) | 48.518103 | 58.265546 | 1.38 | 10 | 13300 | 14411 | 16080 | 0.001 |
| Goose Arm Estuary (gae) | 49.175184 | 57.85118 | NA | 0 | 11000 | 0 | 0 | 0.000 |
| George's Lake (gel) | 48.779225 | 58.103116 | 15.97 | 148 | 12700 | 32152 | 44395 | 0.005 |
| Glyn Mill Inn Pond (gmp) | 48.946415 | 57.939578 | 0.04 | 20 | 11800 | 2772 | 3159 | 0.006 |
| Gravels Pond (grp) | 48.556744 | 58.724693 | 0.16 | 1 | 13600 | 37 | 37 | 0.027 |
| Gull Pond (gup) | 48.539098 | 58.500473 | 0.61 | 29 | 13400 | 6799 | 8017 | 0.004 |
| Hughes Brook Estuary (bbe) | 48.992047 | 57.902362 | NA | 0 | 11800 | 0 | 0 | 0.000 |
| High Elevation Pond (hep) | 49.286856 | 57.725758 | 0.09 | 192 | 10300 | 17286 | - | 0.011 |

| Location | Lat (°N) | Lon (°W) | Lake Area<br>(km<sup>2</sup>) | Elevation<br>(m a.s.l.) | Estimated<br>age (y) | Distance from the ocean |  | Average<br>Gradient |
| --- | --- | --- | --- | --- | --- | --- | --- | --- |
|  |  |  |  |  |  | Straight<br>line (m) | Run of river<br>(m) |  |
| Heron Pond (hpe) | 47.606262 | 57.651891 | NA | 0 | 14500 | 0 | 0 | 0.000 |
| Island Pond (isp) | 48.716389 | 58.122778 | 0.86 | 170 | 12900 | 37192 | 52288 | 0.005 |
| Little Codroy Estuary (lce) | 47.772671 | 59.258507 | NA | 0 | 14300 | 0 | 0 | 0 |
| Long Gull Pond (lgp) | 48.562832 | 58.469474 | 2.66 | 30 | 13400 | 9220 | 13479 | 0.003 |
| Loch Leven (lol) | 48.163905 | 58.870447 | 0.12 | 40 | 14200 | 7561 | 9576 | 0.005 |
| Long Steady (los) | 50.6171 | 57.1149 | 0.39 | 18 | 8500 | 4099 | 5044 | 0.004 |
| Muddy Hole, Flat Bay (mhe) | 48.401727 | 58.58235 | 0.66 | 2 | 13900 | 1334 | 1743 | 0.000 |
| Pond at Mattis Point Road (mpe) | 48.483834 | 58.418696 | 0.27 | 23 | 13400 | 4830 | 6864 | 0.005 |
| North Belburn Pond (nbp) | 50.3772 | 57.5245 | 0.02 | 27 | 8600 | 1297 | 1874 | 0.021 |
| Noel's Pond (nop) | 48.554398 | 58.531816 | 1.03 | 21 | 13400 | 3277 | 4599 | 0.006 |
| Ned's Pond (nsp) | 48.566557 | 58.575538 | 0.24 | 39 | 13400 | 2696 | 3865 | 0.014 |
| Pinchgut Lake (pgl) | 48.8058 | 58.01261 | 3.48 | 189.5 | 12200 | 48758 | 65855 | 0.004 |
| Pond at Main Brook (pmb) | 51.179739 | 56.041474 | 0.87 | 24 | 9000 | 974 | 974 | 0.001 |
| Estuary at Port aux Mal (pme) | 48.65639 | 58.672623 | NA | 0 | 13700 | 0 | 0 | 0.000 |
| Parson's Pond Estuary (ppe) | 50.030008 | 57.710092 | NA | 0 | 8900 | 0 | 0 | 0.000 |
| Rapid Pond (rap) | 48.998821 | 57.702851 | 0.03 | 20 | 11400 | 14940 | 19025 | 0.001 |
| Rocky Harbour Pond (rhp) | 49.571 | 57.903381 | 1.17 | 30 | 9900 | 3697 | 5238 | 0.008 |
| Rocky Pond (rop) | 49.802873 | 57.866725 | 0.22 | 10 | 9300 | 784 | 1256 | 0.013 |
| Sandy Brook (sab) |  |  | 0.06 | 10 | 8700 | 1535 | 1773 | 0.007 |
| Estuary at St George's (sge) | 48.431178 | 58.477677 | NA | 0 | 13500 | 0 | 0 | 0.000 |
| St. Paul's Estuary (spe) | 49.866 | 57.8184 | NA | 0 | 9100 | 0 | 0 | 0.000 |
| Small Pond near Kippens (spk) | 48.553012 | 58.649585 | 0.03 | 29 | 13500 | 852 | 944 | 0.034 |
| Tilt Pond (tip) | 49.672 | 57.9647 | 0.11 | 0 | 9700 | 92 | 95 | 0.000 |
| Two Mile Pond (tmp) | 49.8303 | 57.82067 | 2.26 | 9 | 9200 | 2331 | 4104 | 0.004 |
| Tompkins Pond (top) | 47.798816 | 59.214875 | 0.06 | 13 | 14500 | 8826 | 12009 | 0.002 |
| Trout River Pond (trp) | 49.4598 | 58.1193 | 2.94 | 8 | 10400 | 3443 | 4086 | 0.002 |
| Western Brook Pond (wbp) | 49.7814 | 57.8409 | 23.36 | 22 | 9300 | 5683 | 9713 | 0.004 |
| Wild Cove (wic) | 48.972031 | 57.88755 | NA | 0 | 11800 | 0 | 0 | 0.000 |

We collected stickleback from each location during early-June to mid-July from 2013 to 2019 using 6 mm mesh Gee-type minnow traps placed at depths ranging from 0.5 to 2.0 m for between 5 and 24 hours, or a 10 m long x 2 m deep, 3 mm mesh beach seine. All fish collected were identified to species, counted, and up to 50 stickleback specimens were fixed in 10 % formalin. We released all other species alive at the point of capture. Stickleback samples were transferred to 50% isopropanol for storage after several weeks in 10% formalin.

All fish in each collection were stained with Alizarin Red S (following methods for bone staining outlined in Song and Parenti 1995) to assist with identification of landmarks. Ten to 20 individuals were haphazardly selected form each sample and photographed (left lateral and ventral orientations; using a Canon 5D MarkIII with a 100 mm macro lens or a Nikon D600 with a 90 mm macro lens). All photographs included a scale bar.

We used ImageJ (Schneider et al. 2012) to measure traits related to size (standard length, head length, body depth) and antipredator armor (1^st^ dorsal spine length, 2^nd^ dorsal spine length, pelvic ascending process length, length of each of the three lateral plates fused to the ascending process, pelvic girdle length, left pelvic spine length and lateral plate number; Fig. 2). Landmarks for body shape analysis (Fig. 2) were digitized using tpsUtil (version 1.78) and tpsDig2 (version 2.31; Rohlf 2015).

Statistical analyses were conducted using RStudio environment (version 2021.09.2+382; RStudio Team 2020) for R (version 1.4.2; R Core Team 2021). For the linear variables, we adjusted for allometric effects following the method outlined by Lleonart et al. (2000) with modifications described in Stuart et al. (2017). Size adjusted values for each individual across all linear traits were calculated using the following:

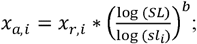

where *x*_*a,i*_ is the size adjusted trait value for individual *i, x*_*r,i*_ is the raw trait value for individual *i, SL* is the average standard length for all individuals measured, *sl*_*i*_ is the standard length for individual *i* and *b* is the slope of the relationship between the log of specific trait against the log of standard length. The slope of the relationship was determined using linear mixed model of each log transformed trait against the log transformed standard length with source population included as a random effect.

We conducted principal components analysis on all size adjusted linear measurements following the methods described in Horst et al. (2020). Briefly, all variables were re-scaled to a mean of 0 and a standard deviation of 1, and principal components were derived from the variance-covariance matrix of the re-scaled variables. We also performed a one-way MANOVA (Quinn and Keough 2002) to examine differences in the size-adjusted, log-transformed trait values between stickleback from either freshwater or marine habitats. Response variables in the MANOVA were population means since we were interested in differences among populations. A significant omnibus test was followed-up with one-way ANOVA comparing marine and freshwater locations for each variable and assessed after Bonferoni correction.

Shape analysis was conducted using MorphoJ (version 1.07a) on unbent landmarks (using landmarks 1, 8 and 21 in tpsUtil; Klingenberg 2011). Landmarks were adjusted for size and orientation using Procrustes fit and the variance-covariance matrix was used to generate new principal component axes based on the adjusted landmark positions. We compared shape PC scores (again using averages for each population as the response variable) between freshwater and marine collection sites (ANOVA, Quinn and Keough 2002).

Finally, we examined correlation between PC scores derived from the size-adjusted linear measurements and the landmark based shape PC scores and attributes of each freshwater system (Table 1).

## Results

We photographed 710 individuals across the 52 sampled locations. One sample was destroyed between taking lateral and ventral photographs (spk) so this population was excluded from the analysis of linear measurements. Additionally, visual inspection of the data identified several outliers. All suspected outliers were inspected and corrected or removed from analyses if appropriate (e.g. broken or missing trait or obscured or missing landmark). Our analysis of size/armor traits included 695 individuals and our analysis of shape included 674 individuals.

### Lateral plates

All marine populations sampled in this study show the typical complete set of armor traits with the exception of a few individuals (Fig. 3). The freshwater locations sampled included low, partial, and complete plate morph populations.

**Figure 3.**
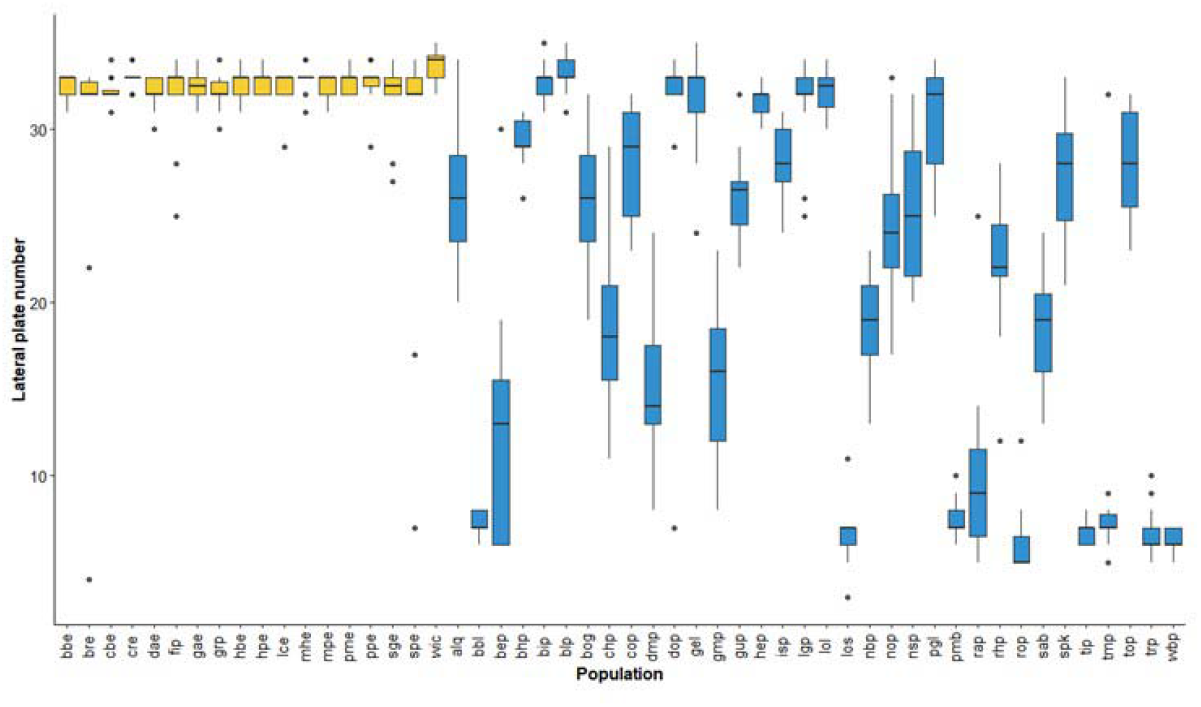
Boxplots of size and armor traits for stickleback sampled at each location. Marine/estuarine sites are indicated with yellow (lighter) shading and freshwater sites are indicated with blue (darker) shading. Labels on the x-axis correspond to the three-letter site codes in Table 1.

### Linear traits

Values for the size-adjusted linear traits are plotted in Fig. 4. Visual inspection of the plots suggests differences in central tendency and dispersion between marine/estuarine populations and freshwater populations. However, one marine/estuarine population (Gravels Pond; grp) does not fall within the bulk of the marine/estuarine populations for spine length traits; spines of individuals in this population appear to be shorter than those for the other marine/estuarine populations. Pearson’s correlation coefficient among traits ranged from 0.56 to 0.93 among the raw linear measurements and between 0.42 and 0.94 for the size-adjusted linear measurements (Table 2; all p-values well below 0.001).

**Table 2.** Pearson correlation coefficient (r) between the variables. Values above the diagonal are for non-size adjusted trait values and values below the diagonal are for size adjusted traits (except for standard length). All p-values <<0.01.

|  | Standard length | Head length | Body depth | Mean lateral plate length | Ascending process length | 1st dorsal spine length | 2nd dorsal spine length | Pelvic girdle length | Pelvic spine length |
| --- | --- | --- | --- | --- | --- | --- | --- | --- | --- |
| Standard length | 1.000 | 0.918 | 0.932 | 0.874 | 0.784 | 0.721 | 0.756 | 0.837 | 0.653 |
| Head length | 0.863 | 1.000 | 0.932 | 0.814 | 0.747 | 0.633 | 0.657 | 0.729 | 0.557 |
| Body depth | 0.887 | 0.827 | 1.000 | 0.918 | 0.862 | 0.721 | 0.755 | 0.817 | 0.666 |
| Mean lateral plate length | 0.813 | 0.723 | 0.878 | 1.000 | 0.847 | 0.759 | 0.780 | 0.840 | 0.661 |
| Ascending process length | 0.695 | 0.642 | 0.812 | 0.792 | 1.000 | 0.747 | 0.757 | 0.817 | 0.614 |
| 1st dorsal spine length | 0.614 | 0.513 | 0.637 | 0.690 | 0.677 | 1.000 | 0.952 | 0.755 | 0.833 |
| 2nd dorsal spine length | 0.663 | 0.524 | 0.664 | 0.703 | 0.681 | 0.941 | 1.000 | 0.794 | 0.850 |
| Pelvic girdle length | 0.762 | 0.604 | 0.734 | 0.772 | 0.756 | 0.685 | 0.724 | 1.000 | 0.633 |
| Pelvic spine length | 0.548 | 0.418 | 0.570 | 0.567 | 0.512 | 0.792 | 0.814 | 0.529 | 1.000 |

**Figure 4.**
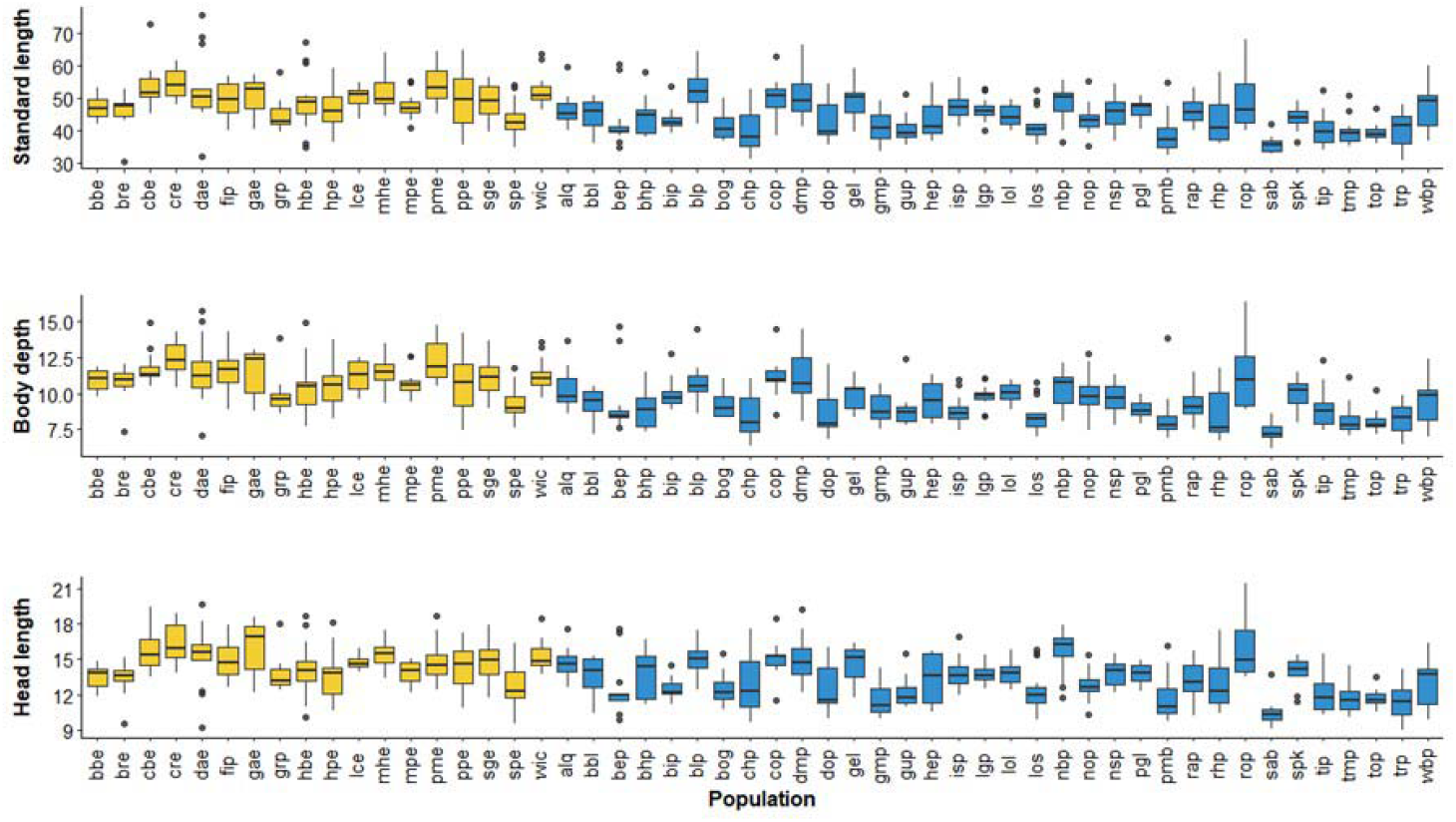
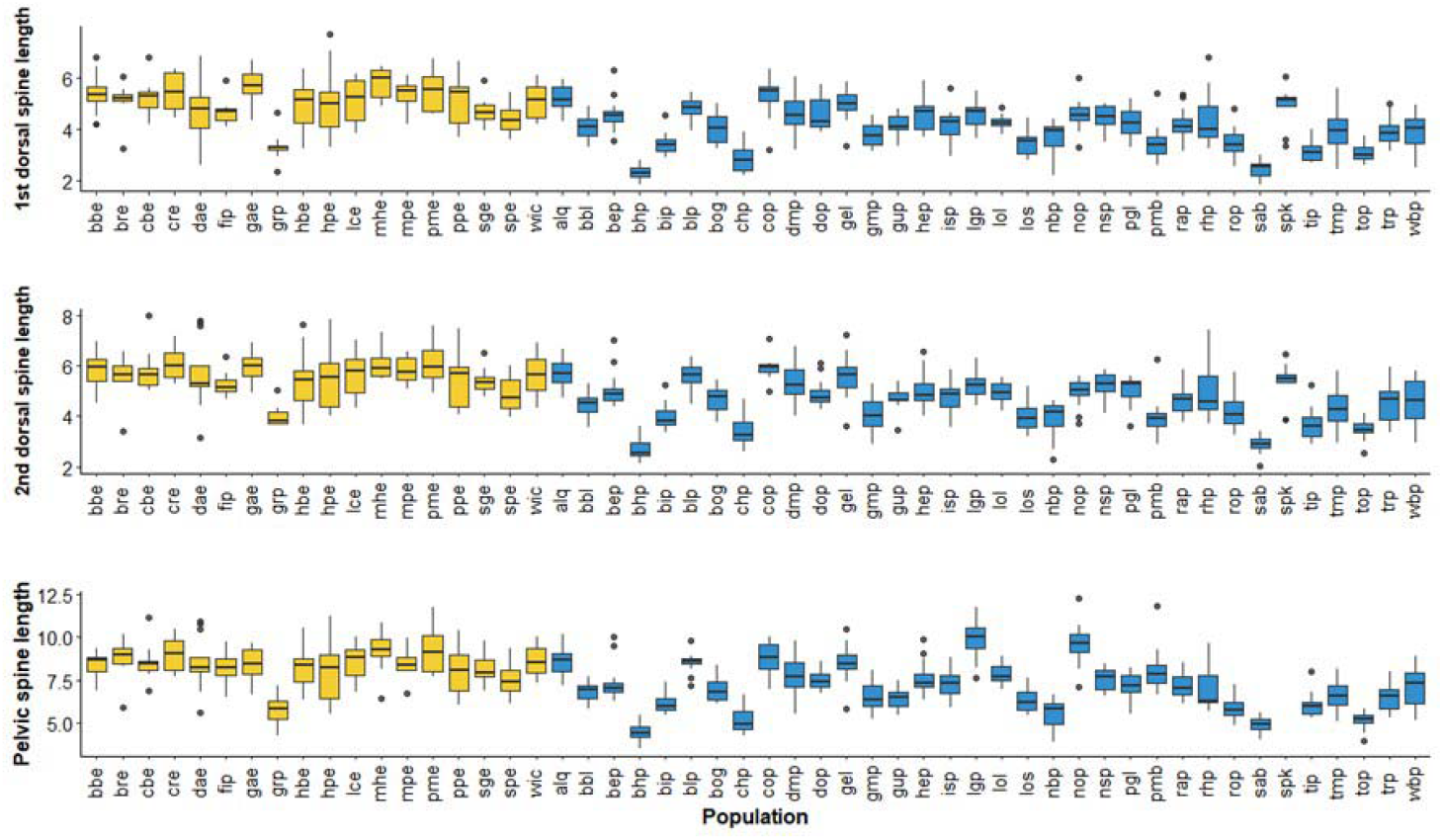
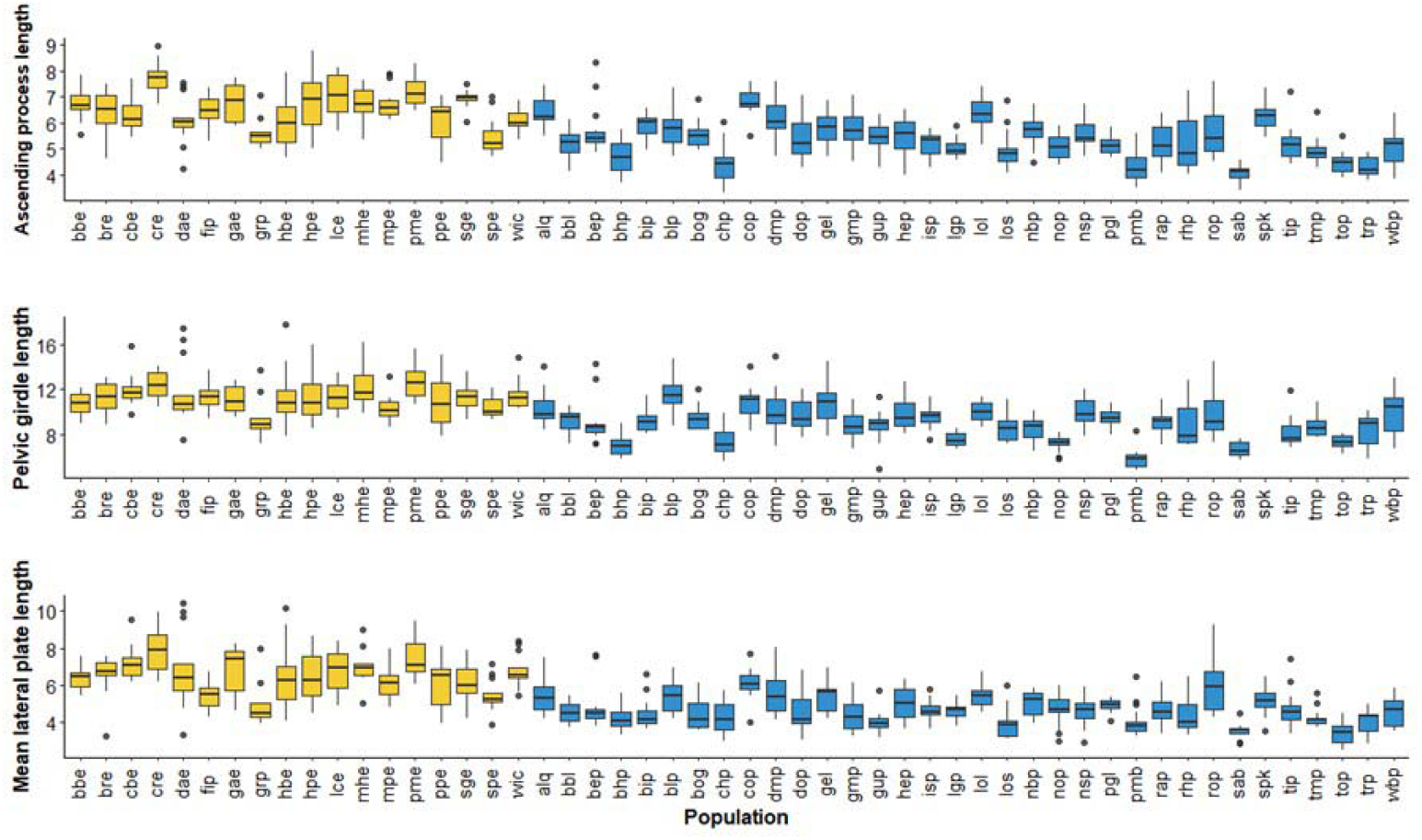
Boxplot of size-adjusted linear traits among populations. Freshwater populations are indicated in blue (darker) and marine/estuarine populations in yellow (lighter).

The first three principal components for the linear variables explained approximately 90% of total variance among individuals (PC1, PC2 and PC3 explained approximately 73%, 12% and 5% respectively). PC1 is positively correlated with all linear measurements (Table 3), PC2 is positively correlated with spine length (both dorsal spines and the pelvic spine), and negatively correlated with head length), and PC3 is negatively correlated with ascending process length and pelvic gridle length and positively correlated with head length and pelvic spine length (Table 3). Marine populations appear to have higher PC1, less variable PC2, and lower PC3 scores than freshwater populations (Fig. 5); marine populations are larger across all traits, freshwater populations are more variable with regard to spine length and head length and freshwater populations appear to have smaller pelvic girdles.

**Table 3.** Factor loadings for each of the size variables for principal components 1-5. The analysis was performed on size adjusted trait values.

|  | PC1 | PC2 | PC3 | PC4 | PC5 |
| --- | --- | --- | --- | --- | --- |
| Standard length | 0.349 | -0.286 | 0.271 | -0.313 | 0.0797 |
| Body depth | 0.356 | -0.271 | 0.0919 | 0.269 | 0.219 |
| Head length | 0.31 | -0.428 | 0.468 | -0.061 | -0.422 |
| 1st dorsal spine | 0.331 | 0.433 | -0.0182 | -0.0162 | -0.475 |
| 2nd dorsal spine | 0.339 | 0.417 | -0.00243 | -0.133 | -0.288 |
| Mean lateral plate length | 0.354 | -0.154 | -0.149 | 0.197 | 0.321 |
| Ascending process length | 0.334 | -0.108 | -0.505 | 0.553 | -0.183 |
| Pelvic spine length | 0.289 | 0.513 | 0.42 | 0.167 | 0.518 |
| Pelvic girdle length | 0.334 | -0.0377 | -0.495 | -0.66 | 0.234 |

**Figure 5.**
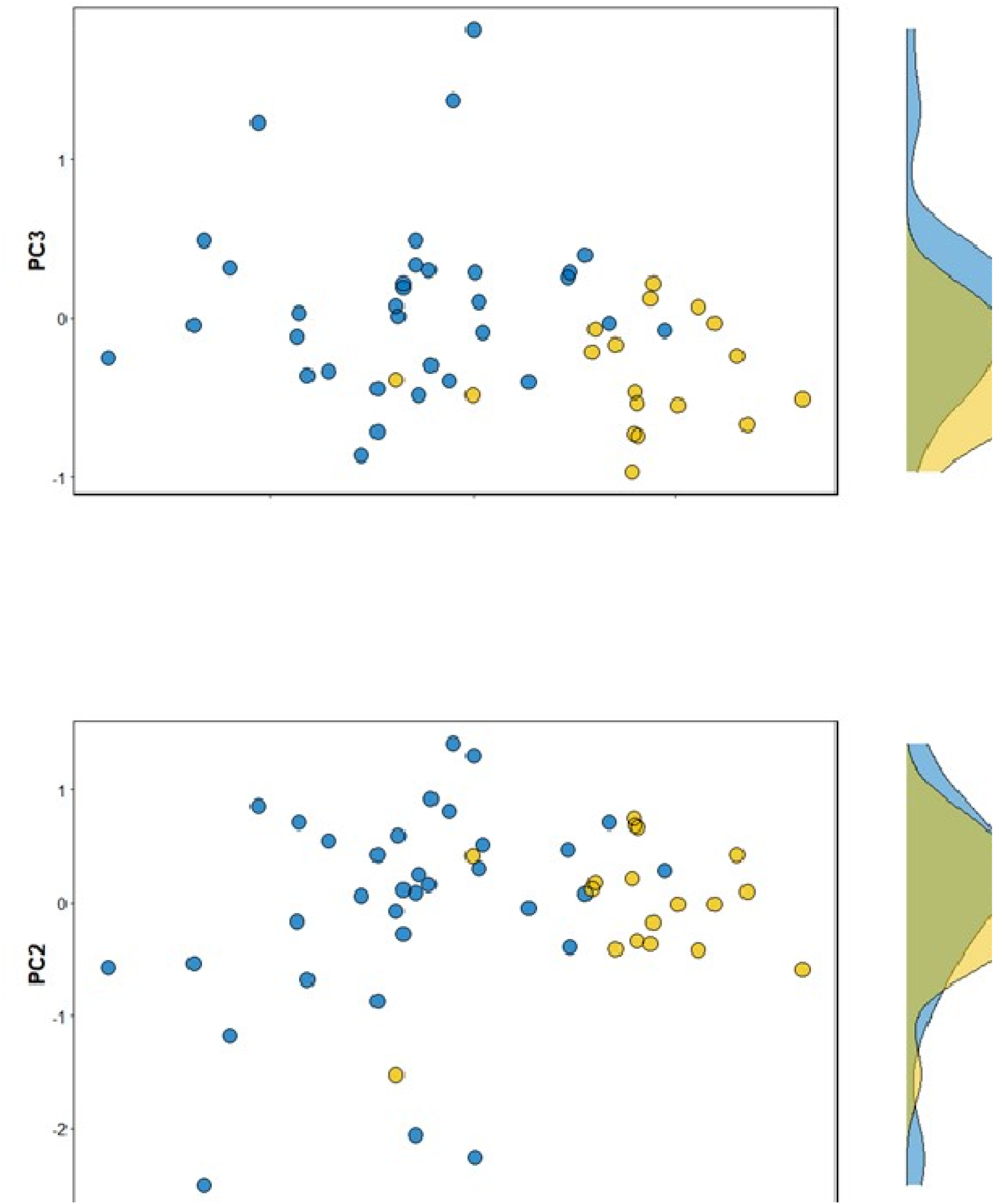
Plot of mean population PC scores and PC score distribution (margins) for linear traits among populations. Freshwater populations are indicated by blue (darker) dots and marine/estuarine populations are yellow (lighter) dots.

MANOVA comparing population mean size-adjusted traits between freshwater and marine sites showed that there is a difference between these types of populations in at least one of the traits (Pillai’s Trace Omnibus F_3, 37_ = 12.889, p<<0.001). All size-adjusted traits were larger for stickleback from marine systems than from freshwater systems (one-way ANOVA for each trait with Bonferroni correction; Table 4; Fig. 6).

**Table 4.** Effect size for habitat (marine versus freshwater) based on one-way ANOVA on log transformed, size-adjusted (except for standard length) values for each variable. Adjusted p value is the Bonferoni corrected p value accounting for multiple tests.

| Variable | Effect size | p value | adjusted p value |
| --- | --- | --- | --- |
| Standard length | 0.0528 | 2.54e- 5 | 2.28e- 4 |
| Body depth | 0.0638 | 7.12e- 8 | 6.41e- 7 |
| Head length | 0.0352 | 1.29e- 3 | 1.16e- 2 |
| 1st dorsal spine length | 0.0988 | 3.88e- 5 | 3.49e- 4 |
| 2nd dorsal spine length | 0.0867 | 9.80e- 5 | 8.82e- 4 |
| Mean lateral plate length | 0.129 | 4.25E-12 | 3.82E-11 |
| Ascending process length | 0.0795 | 6.02e- 7 | 5.42e- 6 |
| Pelvic spine length | 0.0763 | 2.68e- 4 | 2.42e- 3 |
| Pelvic girdle length | 0.0897 | 4.49e- 7 | 4.04e- 6 |

**Figure 6.**
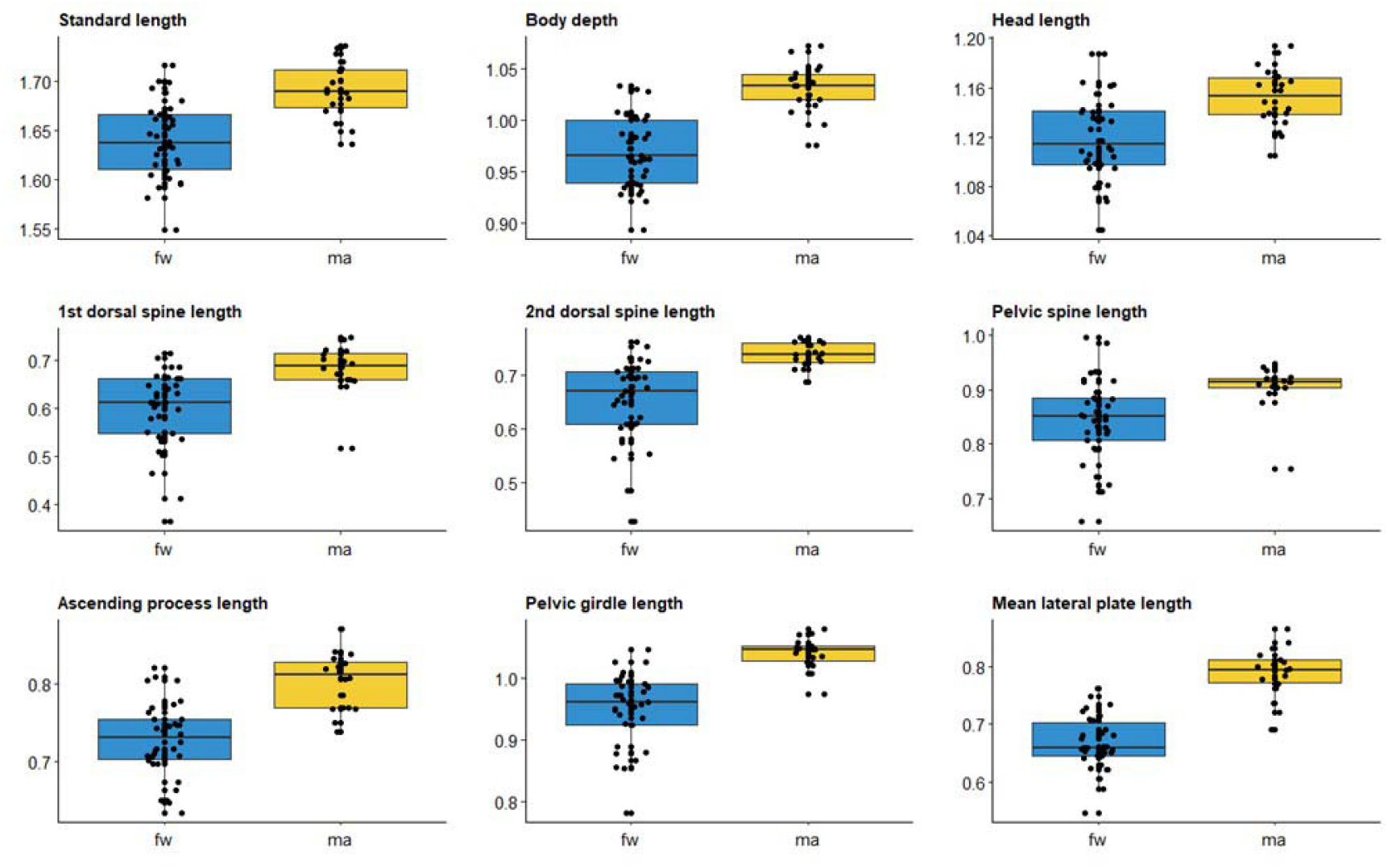
Boxplots of log-transformed, size-adjusted linear traits for marine and freshwater populations of threespine stickleback from western Newfoundland. Individual dots are mean values for each location sampled.

### Body shape

Several principal components were extracted from the landmark data. The first three principal components explain 51% of the total landmark variability (PC1, PC2 and PC3 explain 22%, 18%, and 11% respectively). Shape PC1 is associated with head size and shape with positive values associated with smaller head, eye, and jaw and pectoral fin located forward on the body, whereas negative values associated with larger head, eye and jaw and pectoral fin located caudally (Fig. 7). PC2 is associated with body shape; positive values of PC2 are associated with deeper body and a robust caudal peduncle and negative values with shallow body and thin caudal peduncle (Fig. 7). PC3 is associated with jaw shape; positive values of PC3 are associated with smaller jaw and negative values with larger jaw (Fig. 7).

**Figure 7.**
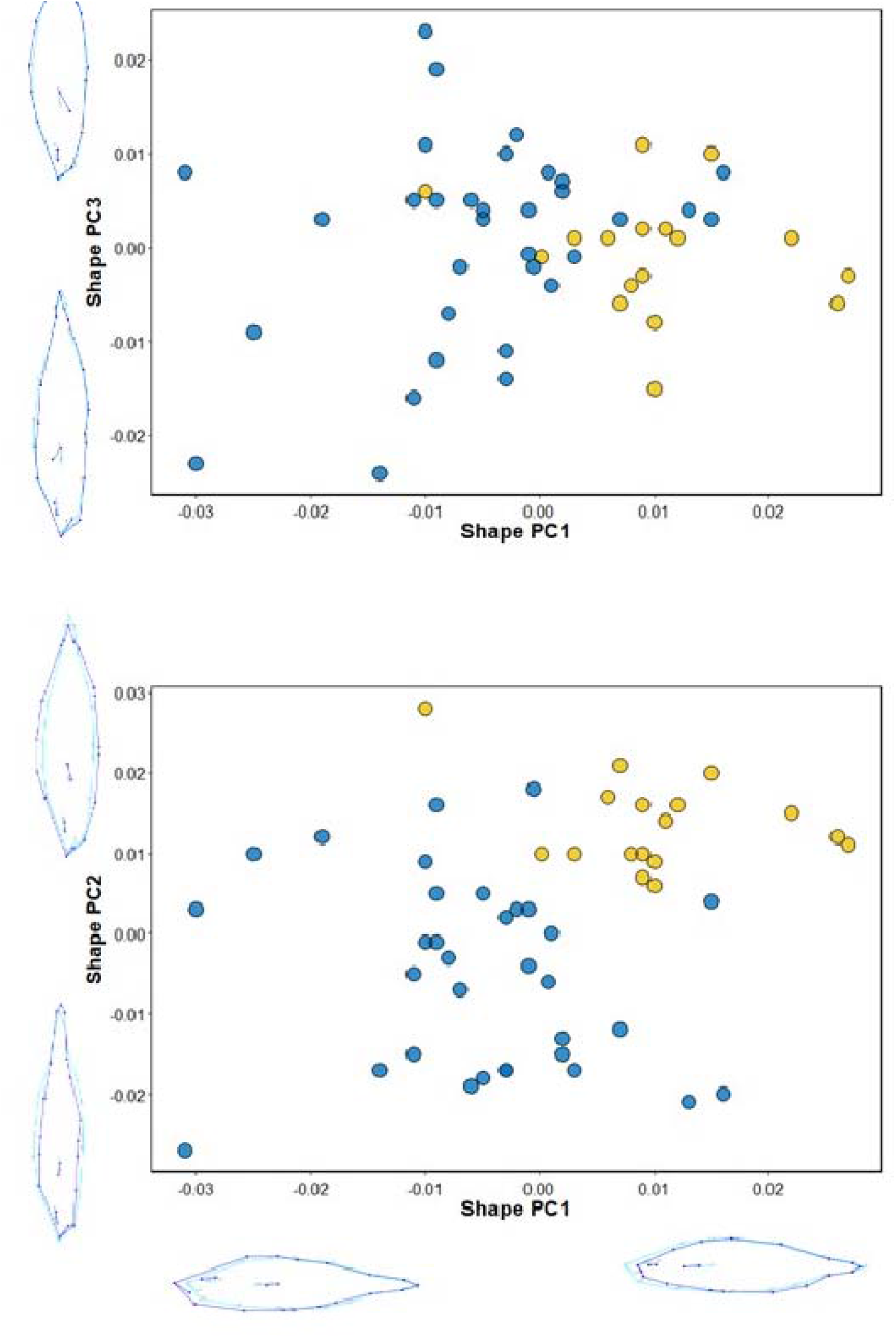
Plot of mean population PC scores and PC score distribution (margins) for shape across populations. Freshwater populations are indicated by blue (darker) dots and marine/estuarine populations are yellow (lighter) dots. Wireframe diagrams are attached to each axis to show the direction of shape change (dark outline) at the ends of each axis relative to the average (light outline).

Marine/estuarine populations tend to score higher on both PC1 and PC2 for shape compared to freshwater populations, but there was no difference between the two types of populations for PC3 (Table 5; Fig. 7). These differences in PC values describe a shorter head, a deeper body and more robust caudal peduncle, with a smaller eye and mouth and more cranially positioned pectoral fin in marine populations compared to fish in freshwater populations.

**Table 5.** Results (F and p) values for one-way ANOVAs comparing PC scores between freshwater and marine/estuarine populations. Numerator df = 1 and denominator df = 48.

|  |  | F | p |
| --- | --- | --- | --- |
| Shape | PC1 | 25.46 | <<0.0001 |
|  | PC2 | 37.45 | <<0.0001 |
|  | PC3 | 0.25 | 0.619 |

### Correlation among LinearPC scores and Lake variables

Lake age was negatively correlated with linear PC 1 and negatively correlated with Shape PC 2 and PC3, so older lakes tend to have larger fish that are shallower bodied. Lake area was positively correlated with Shape PC 1, so larger lakes appear to have fish that have deeper bodies and shorter heads (Table 6).

## Discussion

Replicated adaptive divergence in post-glacial Holarctic fishes provides exceptional opportunities to understand the factors that contribute to the production of new species (Foster et al. 1998; Ehlinger 1999; Schluter 2000a; Ducette et al. 2004; Losos 2010; Magalhaes et al. 2016). Threespine stickleback in freshwater systems demonstrate parallel variation in feeding and antipredator morphology in all regions of their distribution where they have been examined extensively. Frequently, freshwater populations have diverged from the more generalist form of the marine ancestor to specialize on either planktonic or benthic prey. Divergence in feeding morphology among freshwater populations is most extreme when benthic and limnetic forms are sympatric (e.g. Lavin and McPhail 1986; Schluter and McPhail 1992), but can also be observed in allopatric lake populations (Willacker et al. 2010) and adjacent lake and stream populations (Hendry and Taylor 2004, Haenel et al. 2021). Freshwater populations also frequently exhibit reduced antipredator armor, demonstrating reduced number of lateral armor plates, smaller pelvic girdle, and shorter spines (e.g. Bell and Orti 1994).

Haines (2022) reviewed papers examining stickleback morphology of populations from the east coast of Canada. Of 94 freshwater stickleback populations reported across all studies spanning southern New Brunswick north to Baffin Island and Newfoundland west to the western shores of Hudson’s Bay, most populations had a complete set of lateral plates, and less commonly a partial set of lateral plates; very few stickleback populations in the reviewed papers were the low plate form and no population was monomorphic for the low plate form. Only a single low-plated individual—from the Matamek River northwest of Anticosti Island—has previously been reported in eastern Canada outside the Arctic drainage basin (Coad and Power 1974). No papers reported variability in feeding related body shape, likely because no studies have examined variation in these traits from eastern North American populations. Our survey of 52 stickleback populations from the west coast of Newfoundland revealed extensive variation among freshwater populations, in both armor and shape traits, that parallels the variability observed elsewhere in the species’ range.

### Armor and size related trait variability

Freshwater populations consistently experience reduction in lateral plate number following colonization by full-plated marine forms (Bell et al. 2004; Barrett 2010). We found that seven of the 34 freshwater populations examined here maintained the ancestral full-plate form. The remaining populations were either the partial-plate (18 of 34), or low plate (9 of 34) form. Spines (dorsal and pelvic) and pelvic girdle structures also tend to decrease in size in many freshwater populations in Pacific and eastern Atlantic regions (Reimchen 1994). We found freshwater populations that span the range observed elsewhere, from robust spines and pelvic girdles similar to marine populations to markedly reduced spines and pelvic girdles. Additionally, we identified one population from western Newfoundland (not included in this study) that has pelvic reduction/loss and dorsal and lateral plate size reduction (Scott et al. 2022). With regard to antipredator armor, our study suggests that diversification among freshwater populations of east coast North American populations is more common and occurs to a greater degree than has been previously reported.

### Body shape

Freshwater populations of threespine stickleback diverge from the generalist marine ancestral form depending on food sources available and predators present in various lakes (McPhail 1984; Spoljaric and Reimchen 2007; Østbye et al. 2016). Some populations become specialists on benthic prey. Stickleback in these populations tend to have smaller eyes, deeper bodies with shorter heads and terminally oriented mouth compared to populations that specialize on planktonic prey (limnetic form), which tend to have larger eyes, shallower bodies, longer head and upturned mouth (McPhail 1984; Aquirre and Bell 2012; Willacker et al. 2010; Østbye et al. 2016). Freshwater populations of western Newfoundland span a range of body shapes from forms that are similar to the ancestral marine form, characterized by deep body, short head, terminal mouth and small eye to divergent forms that have larger upturned mouth, shallow body, long head and large eye. With regard to body shape, our study shows that stickleback populations of western Newfoundland show variability that is consistent with variability along the benthic-limnetic continuum found elsewhere in the species’ range.

### Conclusions

Despite prevailing assumptions and their relatively recent arrival from Europe, the threespine stickleback populations of the western Atlantic show remarkable phenotypic diversity, even among the sites surveyed here, all within 300 km of each other. Within the time since their colonization of Newfoundland, some of the populations surveyed in this study have developed strongly divergent phenotypes in isolated populations, though it is not clear at whether any of the populations would experience reproductive isolation. Many of these freshwater populations therefore meet at least Hendry et al.’s (2009) conditions for the second state in the four-state process of speciation, ending with irreversible reproductive isolation. Additionally, some of the pond populations are small enough that evolution is likely to occur at a rapid pace, and continued monitoring of these populations, paired with laboratory experimentation to determine extent of reproductive isolation, could yield important insights about the pace and process of speciation in this relatively recent radiation.

While further work is required to establish patterns of divergence within eastern North America, a more detailed history of the threespine stickleback radiation—acquired through a combination of genomic (Fang et al. 2018), palaeogenomic (Kirch et al. 2021), palaeontologic (Benikke 1997), and geologic (Dyke 2004, Laakkonen et al 2021) methods—would be of great benefit in clarifying the evolutionary and biogeographic history of stickleback in the region. Such efforts could allow the western Atlantic stickleback to provide a powerful replicate to studies of parallel evolution and divergence in the radiations of the Pacific and northern Europe, but with increased genetic constraints (Magalhaes et al. 2021) and a narrower temporal window since establishment for the intervention of disruptive stochastic events (e.g., tsunamis and earthquakes (Lescak et al. 2015)). Even within this study, we show consistent variation across the marine-freshwater transition, which is already the subject of considerable work elsewhere. We also include several barachois—estuaries in the Canadian Atlantic region that are separated from the ocean by depositional sandbars—which could offer an interesting counterpoint to the periodically breaching bar-built estuary environments of California (Garcia-Elfring et al. 2021, Wasserman et al. 2021).

The importance of understanding diversification of species in this region extends beyond the stickleback system. As has been argued for the similarly diverse arctic char (*Salvelinus alpinus*), a detailed understanding of the processes leading to speciation and phenotypic diversification below the species level is necessary for anticipating changes to populations and communities resulting from climate change, species introductions, and habitat modification (Reist et al. 2013). In this context, we hope the present study will serve as a basis for future work on the diversity and functional importance of sticklebacks of Newfoundland and Atlantic Canada more broadly.

## Acknowledgements

J. Moriera, M. Graham, D. Budgill, and Will Rauch-Davis assisted with collections and W. Rauch Davis also assisted with image capture and digitization. This research was supported by the Grenfell Campus Research Fund, approved by the Memorial University Animal Care committee, and conducted under collection permits issued by Parks Canada, and Department of Fisheries and Oceans Canada.

## Data availability

data have been uploaded to Borealis and can be accessed at https://doi.org/10.5683/SP3/2ASNRY following publication of this manuscript in a per-reviewed journal.

## Notes

https://doi.org/10.5683/SP3/2ASNRY

### Competing Interest Statement

The authors have declared no competing interest.

